# Eukaryotic recombinases duplicated after divergence from known asgardarchaeal RadA: implications for the evolution of sex during eukaryogenesis

**DOI:** 10.1101/2025.05.23.655837

**Authors:** Lisa Matsuo, Anna Novák Vanclová, Andrew Pomiankowski, Nick Lane, Joel B. Dacks

## Abstract

The origin of meiotic sex was a key milestone in the evolution of the eukaryotic cell. The DNA recombinases Rad51 and DMC1 have been used previously to trace the timing and origins of the meiotic machinery, and warrant revisiting in the face of the recent increased diversity of reported asgardarchaeal taxa. Here we perform comparative genomics and phylogenetic analyses of RadA protein sequences from a broad sampling of eukaryotic and archaeal taxa. We show that even with increased and new sampling, the eukaryotic Rad51 and DMC1 proteins still resolve separately from any archaeal RadA sequences. Taking into account recent evolutionary cell biological discoveries, our data are most consistent with a scenario whereby the asgardarchaeal host cell was evolving cytoskeletal and membrane-protein machinery that was later incorporated into eukaryotic endomembrane systems following the acquisition of mitochondria and the evolution of sex.

## INTRODUCTION

The evolutionary transition from prokaryotic to eukaryotic cellular organization, termed eukaryogenesis (Vosseberg et al. 2024; Donoghue et al. 2023), was arguably the most fundamental shift in cellular history. It is also one of the more highly contested areas of study in evolutionary biology (O’Malley 2010). In particular, determining the order in which eukaryotic traits such as the nucleus, mitochondria, endomembrane systems and cytoskeleton evolved during eukaryogenesis is crucial to understanding the events and forces that initiated the evolution of, and culminated in, the population of organisms that gave rise to all living eukaryotes (Richards et al. 2024), i.e. the Last Eukaryotic Common Ancestor (LECA). One concrete way of addressing eukaryogenesis is to trace the prokaryotic origins of key cellular machinery and assess whether the eukaryotic versions of that machinery are present in modern prokaryotic lineages, and by implication evolved before, during or after the establishment of the endosymbiotic partnership (Richards et al. 2024).

Across the span of theories for eukaryogenesis (Cavalier-Smith 2002; Martin & Müller 1998; Devos 2021; Guglielmini et al. 2022; López-García & Moreira 2023) there is consensus that the process involved the symbiosis of prokaryotic lineages, each one contributing genetic and cellular machinery, that it had independently developed, to the new eukaryotic configuration. While the possibility of additional players cannot be ruled out, it is generally agreed that there were two main prokaryotic contributors (Vosseberg et al. 2024; Donoghue et al. 2023). An alpha proteobacterial lineage gave rise to the mitochondrion, albeit the precise identity of that lineage remains an interesting subject of inquiry (Muñoz-Gómez et al. 2022).

This became an endosymbiotic partner within a lineage of asgardarchaeota, contributing the cytoskeleton, several endomembrane system components, and nuclear house-keeping machinery (Spang et al. 2015; Eme et al. 2023; Akıl & Robinson 2018; Wollweber et al. 2025). The resulting eukaryotic organism was a cellular and genomic chimaera.

The capacity for sexual reproduction is nearly ubiquitous amongst eukaryotic life, with a deeply conserved dance of cell fusion and a two-step meiosis that involves reciprocal exchange across the whole nuclear genome (Bergero et al. 2021; Colnaghi et al. 2020). A two-step meiosis was inferred to be present in the LECA (Dacks & Roger 1999; Ramesh et al. 2005; Schurko et al. 2009) and has long been recognized as a critical step in eukaryogenesis (Bell 1982). The advent of sexual reproduction would have facilitated the evolution of large genomes, compared with organisms that rely on lateral gene transfer (LGT) (Makarova et al. 2005; Koonin et al. 2004; Colnaghi et al. 2020, 2022). In particular, unlike LGT, reciprocal recombination and homologous pairing along chromosomes in meiotic sex enabled the maintenance of far larger numbers of protein-coding genes in the face of deleterious mutation pressure (Colnaghi et al. 2020). It also permitted the expansion of repeat sequences including gene families, introns, and selfish genetic elements, characteristic features of eukaryotic genomes (Colnaghi et al 2022). Notably, the symbiotic partner that became mitochondria did not evolve meiotic recombination, is largely uniparentally inherited and as a consequence is genetically depauperate in the extreme (Radzvilavicius et al. 2017).

One of the most important prokaryotic enzymes involved in genetic recombination is the RecA gene. The RecA protein is essential for DNA repair, homologous recombination and genome stability. RecA has both bacterial and archaeal (RadA) homologues and two paralogues in eukaryotes (Rad51 and DMC1) (Seitz et al. 1998). Rad51 is expressed in both meiotic and mitotic cells, and plays a key role in double-strand-break repair, whereas DMC1 is expressed solely during meiosis and actively promotes strand exchange in homologous recombination (Brown & Bishop 2014). Rad51 and DMC1 are inferred to have archaeal ancestry, being derived from RadA (Seitz et al. 1998), though past analyses have not found any archaea that encode the duplicated paralogues corresponding to Rad51 and DMC1. But there has been a tremendous recent growth in the discovery of archaea, especially asgardarchaea (eg. (Eme et al. 2023)), as well as a large expansion of eukaryotic representation (see (Leka & Wideman 2024)). Therefore, in order to better understand the timing of the duplication that gave rise to the meiosis-specific eukaryotic homologues, we undertook a molecular evolutionary analysis of RadA, Rad51 and DMC1using updated and extended genomic sampling.

## METHODS

### Dataset Assembly

The Comparative Set (TCS), comprising predicted amino acid (protein) sequences from 196 eukaryotic species, was obtained from EukProt (V3) (Leka & Wideman 2024). Non-asgard archaeal sequences were retrieved from Genbank (https://www.ncbi.nlm.nih.gov/genbank/) and JGI (https://genome.jgi.doe.gov/portal/) deposit, while Asgard archaea sequences were collected from Genbank and the dataset published by (Eme et al. 2023). In total, archaeal sequences (Table S1) were collected from four major taxa: Euryarchaeota (70 species), DPANN (30 species), Asgard (32 species), and TACK (24 species).

### Comparative genomics analyses

Hidden Markov Model (HMM) profiles were constructed and employed to identify homologous sequences using HMMer (EDDY 2009) which has greater sensitivity than BLAST (Altschul et al. 1997). Seed queries were used for initial sequence collection to build the respective HMMs: Rad51 (accession number: CAG38796) and DMC1 (accession number: CAG30372.1). These profiles facilitated the detection and identification of Rad51 and DMC1 genes across the assembled eukaryotic and archaeal dataset. For comprehensive comparative genomic analyses, AMOEBAE (Analysis of MOlecular Evolution with BAtch Entry) was utilised (Barlow et al. 2023). This tool supports high-throughput homology searches and provides detailed analytical summaries, enabling efficient characterisation of gene families across diverse taxa and is freely available on GitHUB (https://github.com/laelbarlow/amoebae).

### Phylogenetic analyses

#### Eukaryotic Phylogenetic Tree

Eukaryotic sequences were aligned using MAFFT (Multiple Alignment using Fast Fourier Transform)(v7.520) (Katoh et al. 2019). The aligned sequences were refined with BMGE (Block Mapping and Gathering with Entropy) (Criscuolo & Gribaldo 2010) to remove poorly aligned regions and enhance the quality of the alignment. This resulted in an alignment of 313 positions for 219 taxa which was used to construct a phylogenetic tree (Figure S1) with IQ-TREE2 (Nguyen et al. 2015) employing ModelFinder (automatically selected model: Q.insect+R7) and performing 1000 ultrafast bootstraps and 100 non-parametric bootstraps in parallel to assess branch support.

#### Archaeal Phylogenetic Tree

Asgard archaeal RadA sequences were selected for phylogenetic analyses, along with a smaller set of verified RadA homologues from diverse archaea other than Asgard sourced from the Uniprot database (2025). The initial phylogenetic tree constructed from these sequences exhibited considerable noise due to long branch artifacts. To address this, sequences contributing to long branches were systematically removed, as were RadB sequences misidentified as RadA (Figure S2). The phylogenetic tree (Figure 1) was then rebuilt by IQ-TREE2 with LG+C20+G+F matrix based on the refined set of 85 RadA sequences (61 Asgard and 24 non-asgard) aligned by MAFFT and trimmed by Trimal (Capella-Gutiérrez et al. 2009) to the final length of 286 positions. Analyses using 1000 ultrafast bootstraps and 100 non-parametric bootstraps were run in parallel.

**Figure 1:**
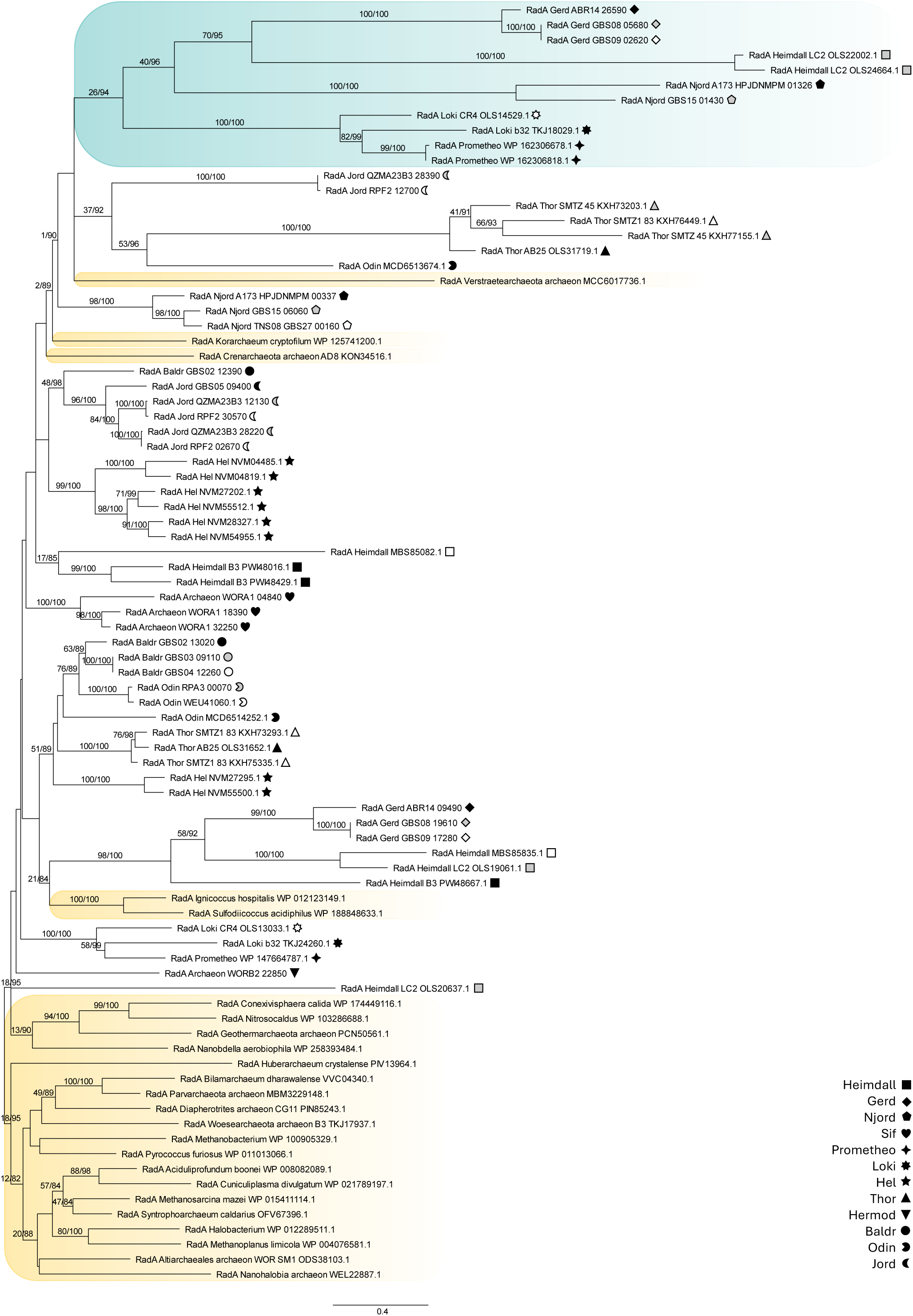
Two RadA clades in a subset of asgardarchaea. This phylogenetic analysis of selected asgardarchaeal RadA homologues reconstructs a moderately supported clade of RadA paralogues (green shade) encompassing Heimdallarchea, Njordarchaea, Gerdarchaea and less robustly Lokiarchaea, distinct from paralogues from the same MAGs in the analysis. Taxon symbols as inset. Support values for 100 NP boostraps and % of 1000 UF boostraps for IQ-TREE analyses are shown at each node supported by greater than 50 NP or 80 UF.

#### Combined datasets

Three additional datasets were prepared using combinations of eukaryotic and archaeal homologues to assess the deeper relationships between RadA paralogs. For these analyses, the eukaryotic sampling was narrowed down to reduce computation time and sampling bias, using a preliminary tree (343 taxa, 300 positions, LG4X matrix, 1000 ultrafast bootstraps; Figure S2) as a reference, resulting in a dataset of 131 DMC1 and Rad51 homologs from 71 organisms. These sequences were then pooled with those from 1) all archaea (Figure S3; tree of 216 taxa based on alignment of 307 positions), 2) Asgard archaea (Figure 2 tree of 192 taxa based on alignment of 306 positions), and 3) a sub-sampling of Asgard archaea representing a presumed eukaryote-adjacent clade/paralogue of RadA (Figure S4; tree of 142 taxa based on alignment of 329 positions). Alignment by MAFFT (Katoh et al. 2019), trimming by Trimal (Capella-Gutiérrez et al. 2009), and tree building by IQ-TREE2 (Nguyen et al. 2015) with LG+C60+G+F matrix and parallel runs using 1000 ultrafast and 100 non-parametric bootstraps, respectively, were used for all three analyses.

**Figure 2:**
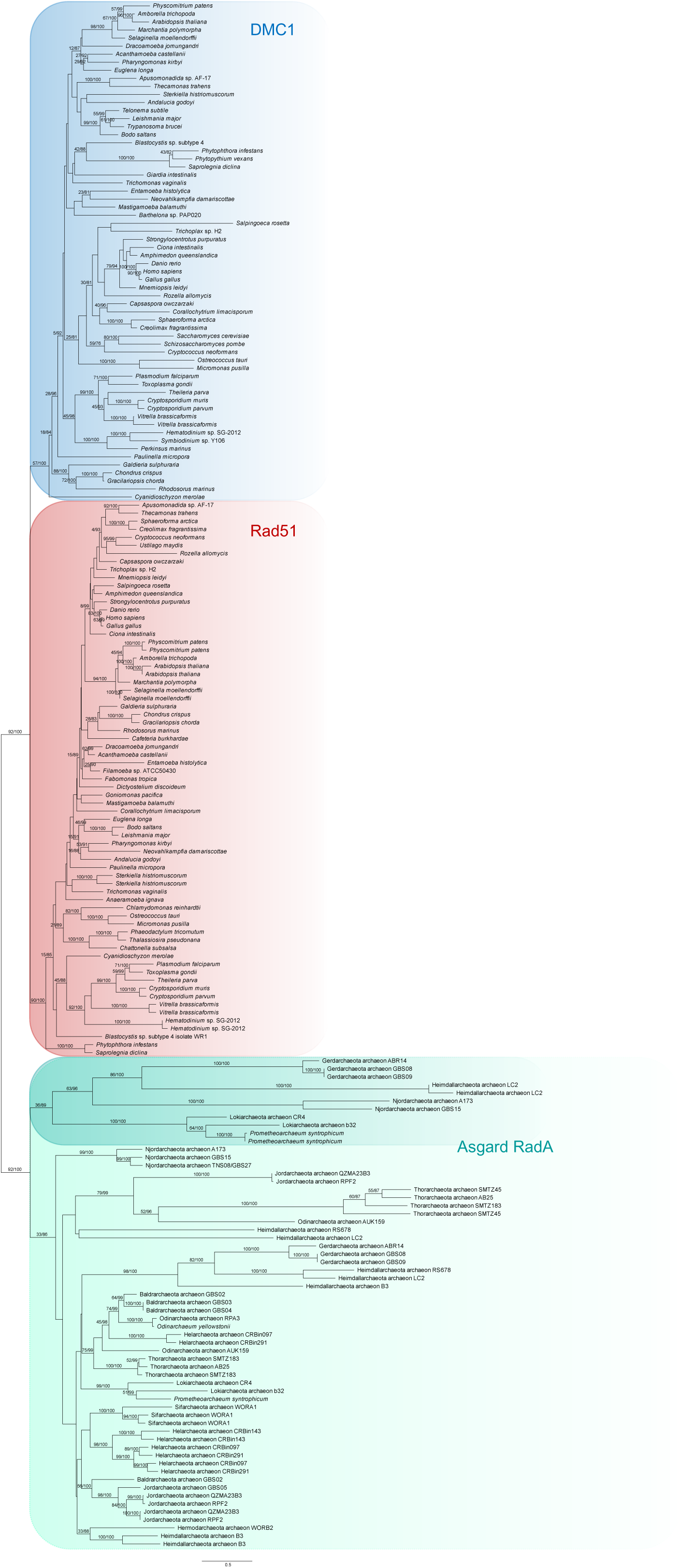
Rad51 and DMC1 form robust eukaryote-specific clades in analysis with all sampled asgardarchaeal RadA paralogues. Support values for 100 NP bootstraps and % of 1000 UF boostraps for IQ-TREE analyses are shown at each node supported by greater than 50 NP or 80 UF.

## RESULTS

In order to address the question of when the recombinases evolved during eukaryogenesis, a comprehensive search was first conducted for Rad51 and DMC1 proteins across a deep and balanced sampling of eukaryotes. Starting from the curated set of eukaryotic genomic and transcriptomic datasets in the EukProt repository (Leka & Wideman 2024), homology searching and validation of ortholog assignment by phylogenetics was used to identify paralogues of both Rad51 and DMC1 in this taxonomically representative sampling of known eukaryotic diversity (Figure S1).

Most notably, we sampled the enigmatic sister-taxon to parabasalids *Anaeramoeba ignava*, the genome of which was recently reported (Jerlström-Hultqvist et al. 2024). Consistent with other reports of unusual meiotic machinery in that taxon (Gallot-Lavallée et al. 2023), there was no DMC1, but Rad51 was found. However, a DMC1 protein was identified in the recently described anaerobic flagellate *Barthelona* sp. PAP020 (Yazaki et al. 2020). While not entirely unexpected, this is an important data point as it is the first molecular evidence in support of a sexual cycle for this protist, a lifestage that has not yet been microscopically observed.

The Rad51 and DMC1 homologues identified were then used to construct HMM profiles which were used as queries for searches into a custom dataset of genomes and Metagenomically Assembled Genomes (MAGs) of archaea (Table S1). This yielded a preliminary dataset, which was merged with the eukaryotic set. Phylogenetic analysis of this larger dataset, allowed us to identify identical, highly divergent, or RadB sequences which had erroneously been drawn in by the sensitive HMM searches (Figure S2)

Using only the archaeal dataset, two clear, moderately supported clades of asgardarchaeal sequences were found (Figure 1), suggesting a duplication of RadA, at the very least in the subgroup uniting Heimdallarchaea and Gerdarchaea, given the relatively robust support for those clades.

Finally, assessment was made of the placement of the eukaryotic recombinases within the prokaryotic RadA homologues. Regardless of whether the analysed datasets included asgardarchaeal and eukaryotic homologues only (Figure 2), non-asgard archaeal outgroups (Figure 3A, Figure S3), or even a restricted dataset (Figure 3B, Figure S4) of the most closely adjacent clade of asgard RadA (as seen in Figure S3), we found the eukaryotic homologues always formed a well-supported monophyletic clade to the exclusion of the archaeal paralogues.

**Figure 3:**
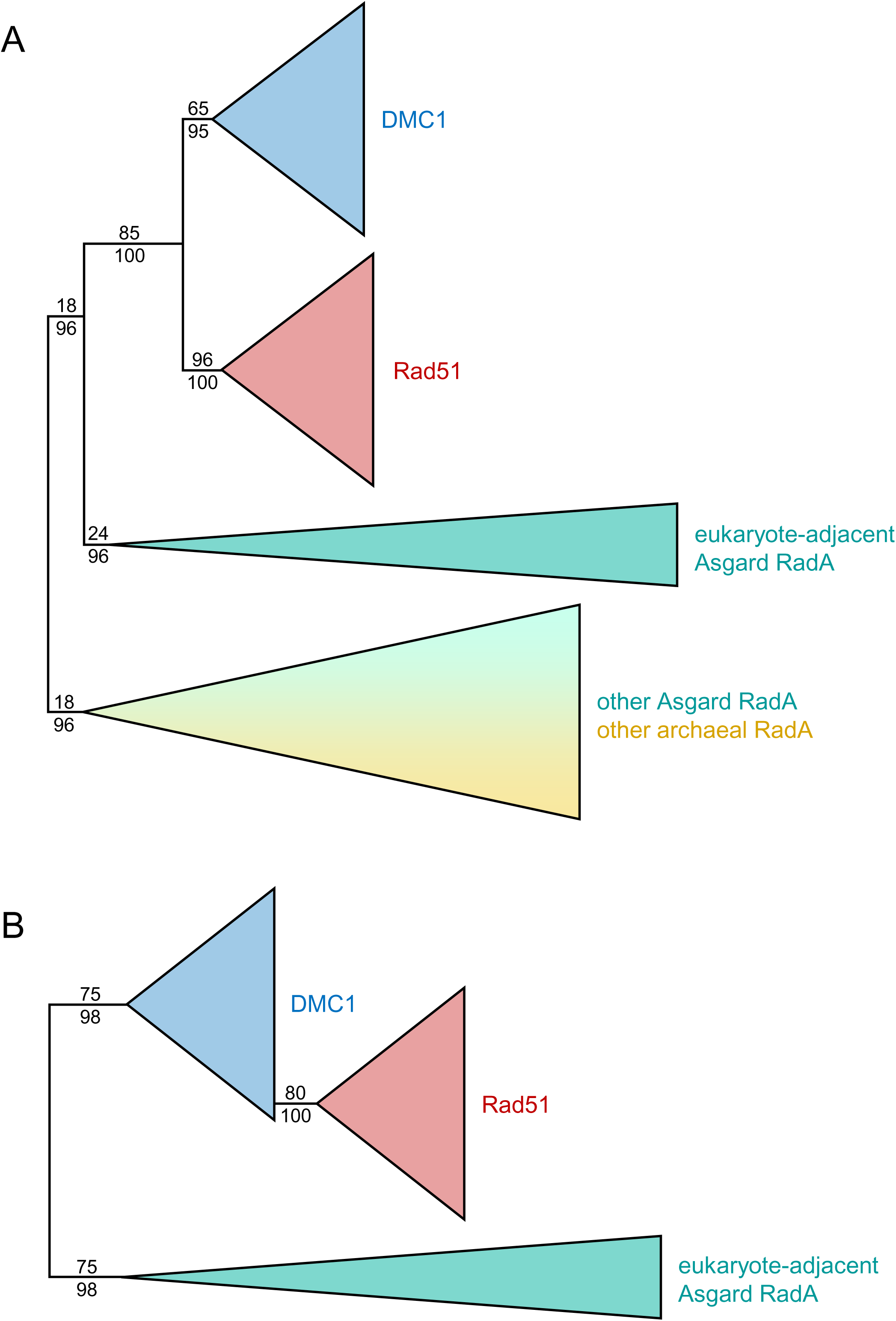
Rad 51 and DMC1 form robust eukaryote-specific clades in analyses with different selections of outgroup taxa. These schematics illustrate the results analyzing the pan-eukaryotic dataset outgroup-rooted by A) the set of RadA paralogues that include asgardarchaeal and non-asgardarchaeal taxa (data shown in full in Figure S3) and B) the set of asgardarchaeal RadA sequences that grouped most closely to eukaryotic paralogs in Figure S3 (data shown in full in Figure S4). Support values for 100 NP bootstraps and % of 1000 UF bootstraps for IQ-TREE analyses are shown only for the clade-defining nodes of interest.

## DISCUSSION

Our phylogenetic investigation shows that the duplication of an ancestral asgardarchaeal RadA gene giving rise to Rad51 and DMC1 took place after the divergence of the eukaryotic lineage from the currently sampled asgardarchaea. If we assume that the Rad51 vs DMC1 duplication informs us about the evolution of meiosis and a sexual cycle, then the data suggest that this eukaryotic hallmark evolved from asgardarchaeal machinery after the divergence of the eukaryotic progenitor from the other asgardarchaeal lineages. Had we observed the duplication in the asgardarchaea, this would have implied meiotic capacity at an earlier stage, prior to the acquisition of the endosymbiont that gave rise to mitochondria.

From the start, it is worth clearly stating limitations to this study. Firstly, these conclusions are only true of the taxa that we sampled, and could be overturned by the discovery of a new asgardarchaeal lineage, particularly one that branches more closely to eukaryotes than do the existing Heimdallarchaeota (Eme et al. 2023), if that organism encoded RadA paralogues that branch with the specific Rad51 and DMC1 clades. Such a lineage has not yet been discovered. Secondly, the assumed scenario of meiosis being tracked by this gene duplication presumes that meiotic function emerged as a eukaryotic feature, which entailed neofunctionalization following duplication. The findings here do not preclude the possibility that some of the meiosis-specific aspects of recombinase function evolved in the pre-duplicated form, as recently suggested elsewhere (Ferlez et al. 2025). Finally, we recognize that we are tracking only a single component of a complicated system, numbering dozens of proteins (Chen & Weir 2024). Understanding the emergence of each of these along the Asgard lineage, or indeed other prokaryotic origins will fill-in the complexity of the story of how meiotic sex evolved during eukaryogenesis. In particular, resolving exactly when eukaryotes acquired fusexins, mostly from the euryarchaea, will be informative for understanding the evolution of cell fusion, which permitted whole genome syngamy (Moi et al. 2022). With these caveats in mind, our data nonetheless lend themselves to some important conclusions about how meiotic sex contributed to eukaryogenesis.

The most straight-forward conclusion is that the duplication of RadA into general and meiosis-specific paralogues, heralding the evolution of sex, did not occur in the asgardarchaeal ancestor of the eukaryotes, but rather after the divergence of the eukaryotic line. This interpretation is consistent with theoretical considerations, which have shown that a large genome, beyond what is seen in bacteria and archaea, cannot be sustained by the genetic exchange facilitated by LGT. This key feature of the eukaryotes is only feasible with recombination between synapsed homologous chromosomes, as in meiosis (Colnaghi et al. 2022, 2020).

This contrasts with features such as the cytoskeleton (Akıl & Robinson 2018; Wollweber et al. 2025) and a number of proteins involved in membrane trafficking and remodelling, notably a well-functioning ESCRT system (Souza et al. 2025) and an asgardarchaeal precursor to the Arf family GTPases (Vargová et al. 2025). These molecular and biochemical signatures have been confirmed in Asgard archaea, even though there is no indication to date that modern asgardarchaea possess extensive internal compartmentalization (Wollweber et al. 2025; Imachi et al. 2025). But the simplest assumption is that these proteins were easily integrated into burgeoning endomembrane systems in early eukaryotes, fashioning their differentiation.

What do the results tell us about the evolution of sex in relation to eukaryogenesis? An obvious hypothesis is that the mitochondrial endosymbiont provided the energy required for a range of complex eukaryotic features, including the expansion of genome size, as seen in LECA (Lane 2020; Lane & Martin 2010). The endosymbiont’s own genome shrank and massively moved into what became the nuclear genome, contributing to its expansion. At the same time, LGT alone has been modelled to be insufficient for the genome expansion necessary for the inferred state of multiple paralogues and extended genomic content in LECA (Lane 2011; Colnaghi et al. 2020, 2022) More work is needed to understand whether the pressure to increase genome size engendered by the presence of the endosymbiont promoted the evolution of meiotic sex or whether meiotic sex preceded and allowed a subsequent increase in genome size. Finally, cytoplasmic fusion and reciprocal recombination likely allowed genetic innovations arising in sub-populations of proto-eukaryotes to be brought together, ultimately giving rise to a relatively homogenous LECA (Záhonová et al. 2025) with a large number of morphological traits not (yet) found in asgardarchaea or any other prokaryotes, including the nucleus, introns and exons, mitosis and meiosis, endomembrane systems and more. Unlike LGT or cloning, reciprocal sex (with some form of cell fusion) can readily explain the accumulation of all these morphological traits in LECA (Lane 2011).

Overall, the data are consistent with some of the most significant genetic and biochemical capabilities underlying eukaryotic cellular configuration (eg. cytoskeleton, membrane-trafficking proteins) evolving in the asgardarchaeal ancestor to eukaryotes, before substantive cellular leaps took place. Powered by the acquisition of a symbiotic partner that eventually became the mitochondrion, sex enabled the evolution of a LECA with a large genome and unprecedented cellular complexity.

## Supporting information

Table of sampled archael genomes and MAGs

## Acknowledgements

We wish to thank Dr. Kristina Zahonova for technical advice, particularly for selection of archael genomes to sample and Dr. Dayana Salas-Leiva for helpful discussion on the evolution of sex, and particularly regarding the meiotic machinery encoded in Anaeramoeba. We also thank the members of the Lane, Pomiankowski and Dacks Labs for technical and collegial support. LM was funded by a Genetics Society Summer Studentship. POM and NL are supported by funding from the Biotechnology and Biological Sciences Research Council (BB/V003542/1). POM is supported by funding from the Engineering and Physical Sciences Research Council (EP/F500351/1, EP/I017909/1) and Natural Environment Research Council (NE/X009734/1). NL is supported by funding from the Bill & Melinda Gates Foundation (<u>inv-064683</u>). Work in the Dacks lab is supported by grants from the Natural Sciences and Engineering Research Council of Canada (RES0043758 and RES0046091).

## Author Contributions

JBD, NL & AP conceived the idea behind the research, LM, ANV and JBD collected the data, carried out the analysis and led the writing of the manuscript. All authors contributed critically to the drafts and gave final approval for publication.

## Data Accessibility Statement

Data and data description are available from https://figshare.com/articles/dataset/Sequence_datasets_alignments_and_phylogenetic_trees/29109308?file=54675668

---

**Figure S1:**
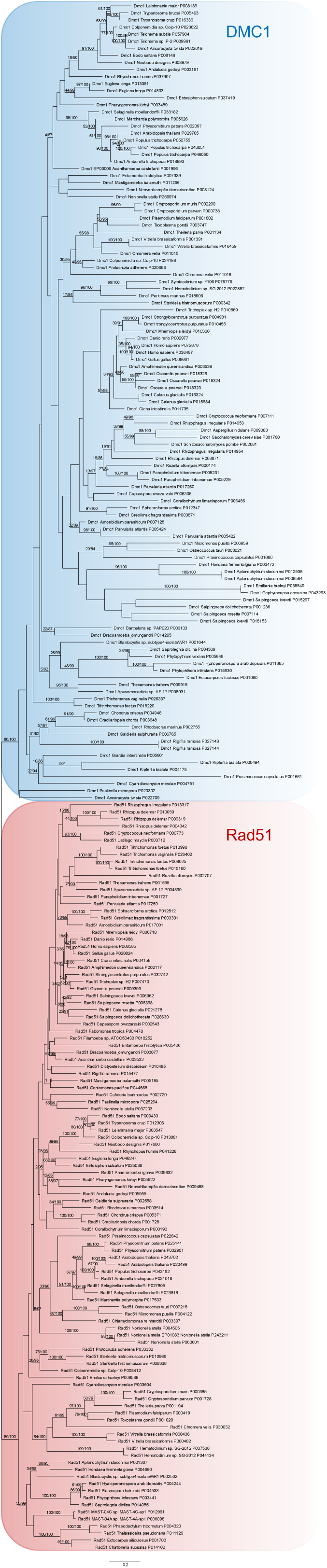
Rad51 and DMC1 are robustly resolved into separate clades encompassing the breadth of eukaryotes. This phylogenetic analysis sampled a wide range of eukaryotic taxa and resolved paralogues into clades of Rad51 (red shading) and DMC1 (blue shading). Notably a DMC1 paralogue was identified in the Barthelona PAP20 genome, a basally branching representative of the Fornicata lineage, for which a sexual cycle has not yet been reported. Support values for 100 NP boostraps and % of 1000 UF boostraps for IQ-TREE analyses are shown at each node supported by greater than 50 NP or 80 UF.

**Figure S2:**
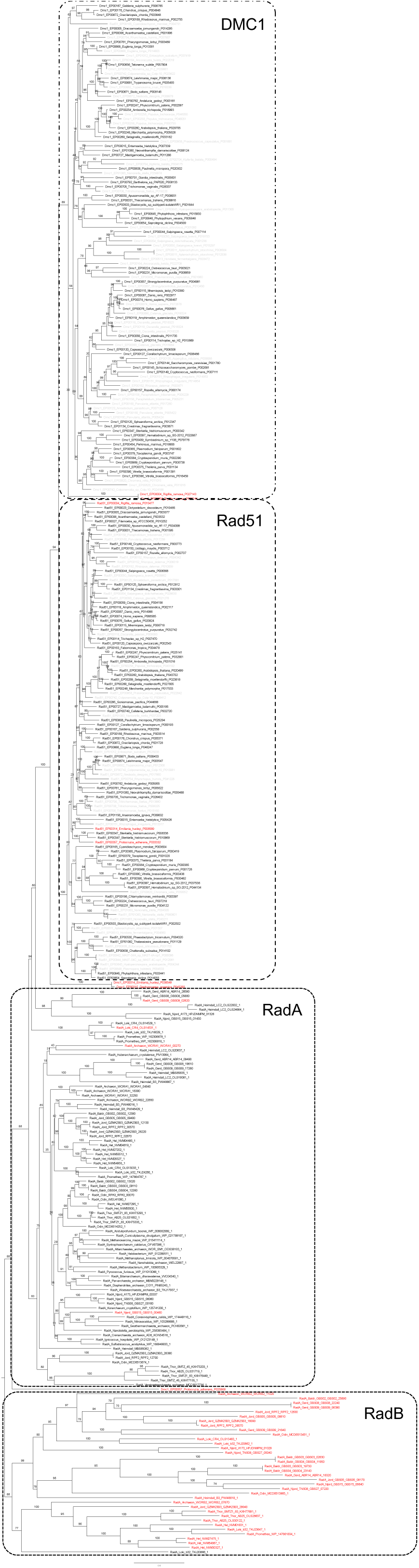
Analysis of all sampled archael Rad protein sequences. This analysis identified identical or highly divergent sequences, as well as RadB outgroup sequences erroneously retrieved by the highly sensitive homology searching approach. Support values for % of 1000 UF boostraps from IQ-TREE analysis and relevant clades are denoted by dashed-line boxes.

**Figure S3:**
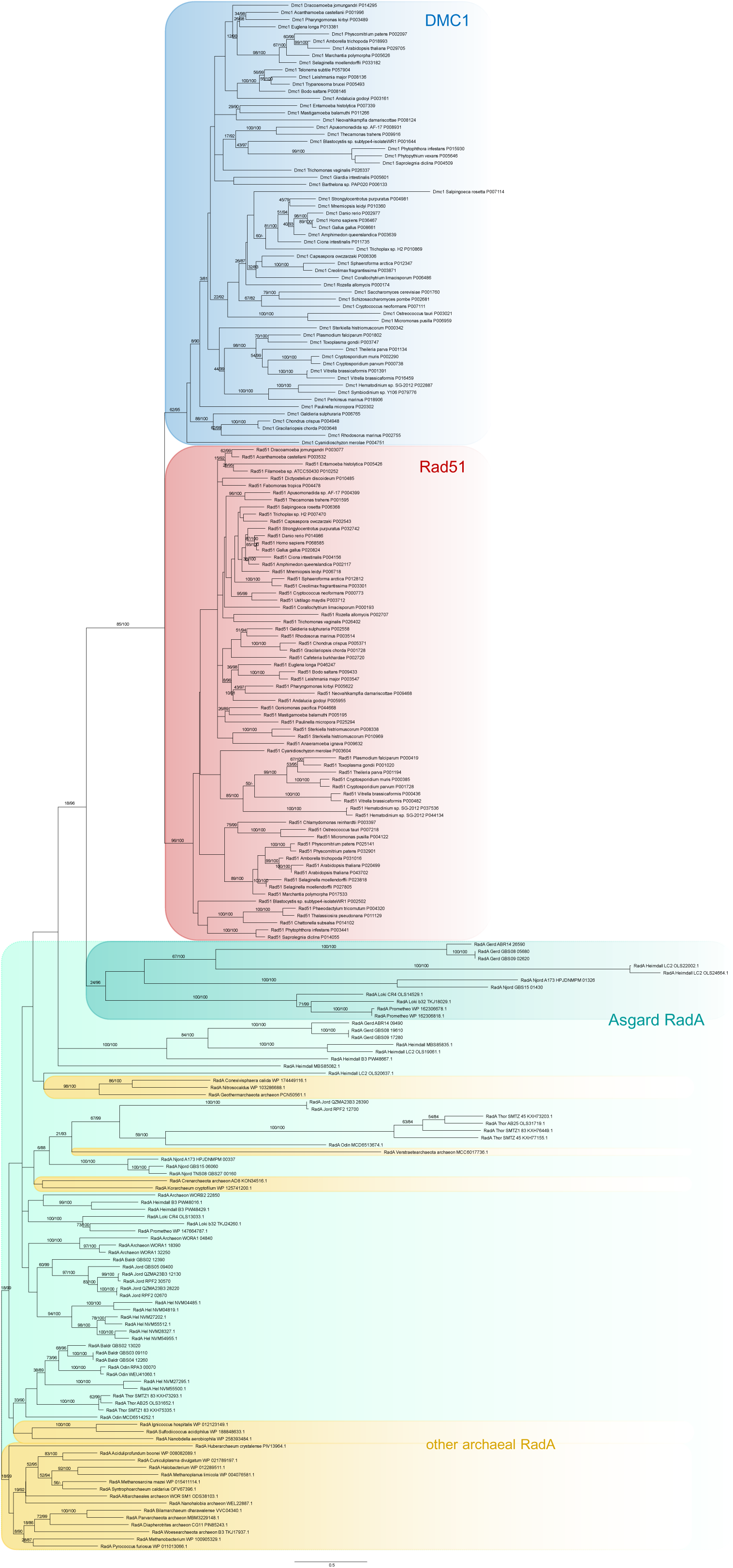
Rad51 and DMC1 form robust eukaryote-specific clades in the presence of asgardarchaeal and non-asgardarchaeal RadA outgroups. Support values for 100 NP bootstraps and % of 1000 UF bootstraps for IQ-TREE analyses are shown at each node supported by greater than 50 NP or 80 UF.

**Figure S4:**
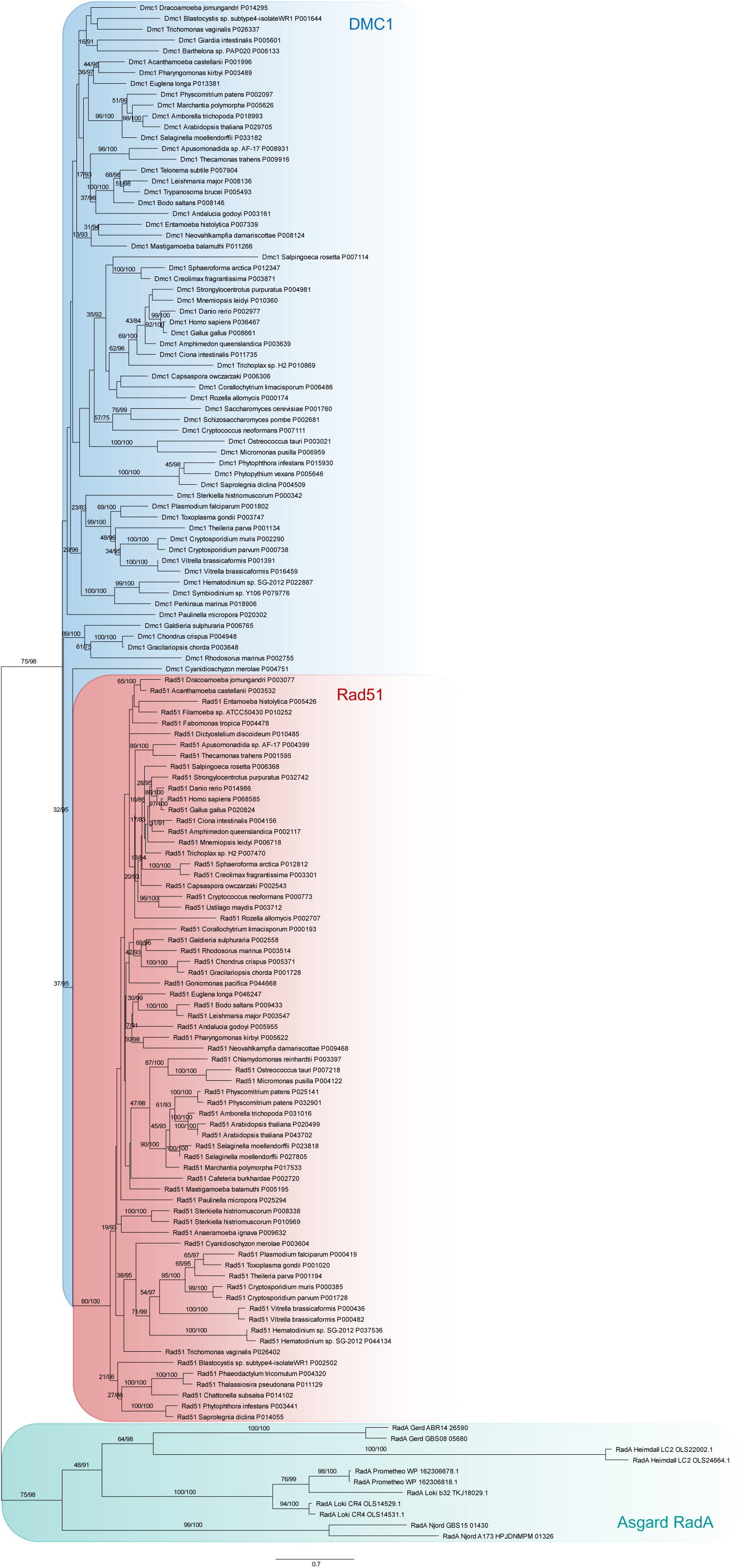
Rad51 and DMC1 duplicated after the divergence from asgardarchaeal RadA. This analysis demonstrates that the eukaryotic Rad51 (red shading) and DMC1 (blue shading) sequences are robustly separated from asgardarchaeal RadA homologues, in this case using only those sequences that resolved closest to the eukaryotic paralogues in an analysis with all archaeal RadA sequences sampled (Figure S3). DMC1 appears paraphyletic in this analysis, in contrast to all other analyses that we performed. Support values for 100 NP bootstraps and % of 1000 UF bootstraps for IQ-TREE analyses are shown at each node supported by greater than 50 NP or 80 UF.

## REFERENCES

Akıl C, Robinson RC. 2018. Genomes of Asgard archaea encode profilins that regulate actin. Nature. 562:439–443. doi: 10.1038/s41586-018-0548-6.

Altschul SF et al. 1997. Gapped BLAST and PSI-BLAST:a new generation of protein database search programs. Nucleic Acids Res. 25:3389–3402. doi: 10.1093/nar/25.17.3389.

Barlow LD et al. 2023. Comparative Genomics for Evolutionary Cell Biology Using AMOEBAE: Understanding the Golgi and Beyond. Methods Mol. Biol. 2557:431–452. doi: 10.1007/978-1-0716-2639-9_26.

Bell G. 1982. The Masterpiece of Nature: The Evolution and Genetics of Sexuality. University of California Press.: Berkeley, CA.

Bergero R et al. 2021. Meiosis and beyond - understanding the mechanistic and evolutionary processes shaping the germline genome. Biol. Rev. Camb. Philos. Soc. 96:822–841. doi: 10.1111/brv.12680.

Brown MS, Bishop DK. 2014. DNA strand exchange and RecA homologs in meiosis. Cold Spring Harb. Perspect. Biol. 7:a016659. doi: 10.1101/cshperspect.a016659.

Capella-Gutiérrez S, Silla-Martínez JM, Gabaldón T. 2009. trimAl: a tool for automated alignment trimming in large-scale phylogenetic analyses. Bioinformatics. 25:1972–1973. doi: 10.1093/bioinformatics/btp348.

Cavalier-Smith T. 2002. The neomuran origin of archaebacteria, the negibacterial root of the universal tree and bacterial megaclassification. Int. J. Syst. Evol. Microbiol. 52:7–76. doi: 10.1099/00207713-52-1-7.

Chen L, Weir JR. 2024. The molecular machinery of meiotic recombination. Biochem. Soc. Trans. 52:379–393. doi: 10.1042/BST20230712.

Colnaghi M, Lane N, Pomiankowski A. 2020. Genome expansion in early eukaryotes drove the transition from lateral gene transfer to meiotic sex. Elife. 9. doi: 10.7554/eLife.58873.

Colnaghi M, Lane N, Pomiankowski A. 2022. Repeat sequences limit the effectiveness of lateral gene transfer and favored the evolution of meiotic sex in early eukaryotes. Proc. Natl. Acad. Sci. U. S. A. 119:e2205041119. doi: 10.1073/pnas.2205041119.

Criscuolo A, Gribaldo S. 2010. BMGE (Block Mapping and Gathering with Entropy): a new software for selection of phylogenetic informative regions from multiple sequence alignments. BMC Evol. Biol. 10:210. doi: 10.1186/1471-2148-10-210.

Dacks J, Roger AJ. 1999. The first sexual lineage and the relevance of facultative sex. J. Mol. Evol. 48:779–783. doi: 10.1007/PL00013156.

Devos DP. 2021. Reconciling Asgardarchaeota Phylogenetic Proximity to Eukaryotes and Planctomycetes Cellular Features in the Evolution of Life. Mol. Biol. Evol. 38:3531–3542. doi: 10.1093/molbev/msab186.

Donoghue PCJ et al. 2023. Defining eukaryotes to dissect eukaryogenesis. Curr. Biol. 33:R919–R929. doi: 10.1016/j.cub.2023.07.048.

Eddy SR. 2009. A NEW GENERATION OF HOMOLOGY SEARCH TOOLS BASED ON PROBABILISTIC INFERENCE. In: Genome Informatics 2009. doi: 10.1142/9781848165632_0019.

Eme L et al. 2023. Inference and reconstruction of the heimdallarchaeial ancestry of eukaryotes. Nature. 618:992–999. doi: 10.1038/s41586-023-06186-2.

Ferlez B, Huang PT, Hu J, Brooks Crickard J. 2025. Evolution of Eukaryotic Specific DNA Binding Sites in Asgard Archaeal RecA Recombinases. bioRxiv. doi: 10.1101/2025.02.19.639135.

Gallot-Lavallée L et al. 2023. Massive intein content in *Anaeramoeba* reveals aspects of intein mobility in eukaryotes. Proc. Natl. Acad. Sci. 120:e2306381120. doi: 10.1073/pnas.2306381120.

Guglielmini J et al. 2022. Viral origin of eukaryotic type IIA DNA topoisomerases. Virus Evol. 8:veac097. doi: 10.1093/ve/veac097.

Hickey DA. 1993. Molecular symbionts and the evolution of sex. J. Hered. 84:410–414. doi: 10.1093/oxfordjournals.jhered.a111363.

Imachi H et al. 2025. Eukaryotes’ closest relatives are internally simple syntrophic archaea. bioRxiv. doi: 10.1101/2025.02.26.640444.

Jerlström-Hultqvist J et al. 2024. A unique symbiosome in an anaerobic single-celled eukaryote. Nat. Commun. 15:9726. doi: 10.1038/s41467-024-54102-7.

Katoh K, Rozewicki J, Yamada KD. 2019. MAFFT online service: multiple sequence alignment, interactive sequence choice and visualization. Brief. Bioinform. 20:1160–1166. doi: 10.1093/bib/bbx108.

Koonin E V et al. 2004. A comprehensive evolutionary classification of proteins encoded in complete eukaryotic genomes. Genome Biol. 5:R7. doi: 10.1186/gb-2004-5-2-r7.

Lane N. 2011. Energetics and genetics across the prokaryote-eukaryote divide. Biol. Direct. 6:35. doi: 10.1186/1745-6150-6-35.

Lane N. 2020. How energy flow shapes cell evolution. Curr. Biol. 30:R471–R476. doi: 10.1016/j.cub.2020.03.055.

Lane N, Martin W. 2010. The energetics of genome complexity. Nature. 467:929–934. doi: 10.1038/nature09486.

Leka KP, Wideman JG. 2024. An introduction to comparative genomics, EukProt, and the reciprocal best hit (RBH) method for bench biologists: Ancestral phosphorylation of Tom22 in eukaryotes as a case study. Methods Enzymol. 707:209–234. doi: 10.1016/bs.mie.2024.07.036.

López-García P, Moreira D. 2023. The symbiotic origin of the eukaryotic cell. C. R. Biol. 346:55–73. doi: 10.5802/crbiol.118.

Makarova KS, Wolf YI, Mekhedov SL, Mirkin BG, Koonin E V. 2005. Ancestral paralogs and pseudoparalogs and their role in the emergence of the eukaryotic cell. Nucleic Acids Res. 33:4626–4638. doi: 10.1093/nar/gki775.

Martin W, Müller M. 1998. The hydrogen hypothesis for the first eukaryote. Nature. 392:37–41. doi: 10.1038/32096.

Moi D et al. 2022. Discovery of archaeal fusexins homologous to eukaryotic HAP2/GCS1 gamete fusion proteins. Nat. Commun. 13:3880. doi: 10.1038/s41467-022-31564-1.

Muñoz-Gómez SA et al. 2022. Site-and-branch-heterogeneous analyses of an expanded dataset favour mitochondria as sister to known Alphaproteobacteria. Nat. Ecol. Evol. 6:253–262. doi: 10.1038/s41559-021-01638-2.

Nguyen L-T, Schmidt HA, von Haeseler A, Minh BQ. 2015. IQ-TREE: A Fast and Effective Stochastic Algorithm for Estimating Maximum-Likelihood Phylogenies. Mol. Biol. Evol. doi: 10.1093/molbev/msu300.

O’Malley MA. 2010. The first eukaryote cell: an unfinished history of contestation. Stud. Hist. Philos. Biol. Biomed. Sci. 41:212–224. doi: 10.1016/j.shpsc.2010.07.010.

Radzvilavicius AL, Lane N, Pomiankowski A. 2017. Sexual conflict explains the extraordinary diversity of mechanisms regulating mitochondrial inheritance. BMC Biol. 15:94. doi: 10.1186/s12915-017-0437-8.

Ramesh MA, Malik S-B, Logsdon JMJ. 2005. A phylogenomic inventory of meiotic genes; evidence for sex in Giardia and an early eukaryotic origin of meiosis. Curr. Biol. 15:185–191. doi: 10.1016/j.cub.2005.01.003.

Richards TA et al. 2024. Reconstructing the last common ancestor of all eukaryotes. PLoS Biol. 22:e3002917. doi: 10.1371/journal.pbio.3002917.

Schurko AM, Neiman M, Logsdon JMJ. 2009. Signs of sex: what we know and how we know it. Trends Ecol. Evol. 24:208–217. doi: 10.1016/j.tree.2008.11.010.

Seitz EM, Brockman JP, Sandler SJ, Clark AJ, Kowalczykowski SC. 1998. RadA protein is an archaeal RecA protein homolog that catalyzes DNA strand exchange. Genes Dev. 12:1248–1253. doi: 10.1101/gad.12.9.1248.

Souza DP et al. 2025. Asgard archaea reveal the conserved principles of ESCRT-III membrane remodeling. Sci. Adv. 11:eads5255. doi: 10.1126/sciadv.ads5255.

Spang A et al. 2015. Complex archaea that bridge the gap between prokaryotes and eukaryotes. Nature. doi: 10.1038/nature14447.

Vargová R et al. 2025. The Asgard archaeal origins of Arf family GTPases involved in eukaryotic organelle dynamics. Nat. Microbiol. 10:495–508. doi: 10.1038/s41564-024-01904-6.

Vosseberg J et al. 2024. The emerging view on the origin and early evolution of eukaryotic cells. Nature. 633:295–305. doi: 10.1038/s41586-024-07677-6.

Wollweber F et al. 2025. Microtubules in Asgard archaea. Cell. doi: 10.1016/j.cell.2025.02.027.

Yazaki E et al. 2020. Barthelonids represent a deep-branching metamonad clade with mitochondrion-related organelles predicted to generate no ATP. Proceedings. Biol. Sci. 287:20201538. doi: 10.1098/rspb.2020.1538.

Záhonová K, Lukeš J, Dacks JB. 2025. Diplonemid protists possess exotic endomembrane machinery, impacting models of membrane trafficking in modern and ancient eukaryotes. Curr. Biol. doi: 10.1016/j.cub.2025.02.032.

2025. UniProt: the Universal Protein Knowledgebase in 2025. Nucleic Acids Res. 53:D609–D617. doi: 10.1093/nar/gkae1010.

